# Multi-trait ensemble genomic prediction and simulations of recurrent selection highlight importance of complex trait genetic architecture in long-term genetic gains in wheat

**DOI:** 10.1101/2022.11.08.515457

**Authors:** Nick Fradgley, Keith A. Gardner, Alison R. Bentley, Phil Howell, Ian J. Mackay, Michael F. Scott, Richard Mott, James Cockram

## Abstract

Cereal crop breeders have achieved considerable genetic gain in genetically complex traits, such as grain yield, while maintaining genetic diversity. However, focus on selection for yield has negatively impacted other important traits. To better understand selection within a breeding context, and how it might be optimised, we analysed genotypic and phenotypic data from a diverse, 16-founder wheat multi-parent advanced generation inter-cross (MAGIC) population.

Compared to single-trait models, multi-trait ensemble genomic prediction models increased prediction accuracy for almost 90% of traits, improving grain yield prediction accuracy by 3-52%. For complex traits, non-parametric models (Random Forest) also outperformed simplified, additive models (LASSO), increasing grain yield prediction accuracy by 10-36%. Simulations of recurrent genomic selection then showed that sustained greater forward prediction accuracy optimised long-term genetic gains.

Simulations of selection on grain yield found indirect responses in related traits, which involved optimisation of antagonistic trait relationships. We found multi-trait selection indices could be used to optimise undesirable relationships, such as the trade-off between grain yield and protein content, or combine traits of interest, such as yield and weed competitive ability.

Simulations of phenotypic selection found that including Random Forest rather than LASSO genetic models, and multi-trait rather than single-trait models as the true genetic model, accelerated and extended long-term genetic gain whilst maintaining genetic diversity. These results suggest important roles of pleiotropy and epistasis in the wider context of wheat breeding programmes and provide insights into mechanisms for continued genetic gain in a limited genepool and optimisation of multiple traits for crop improvement.

## 1. Introduction

Classical plant breeding aims to achieve continuous genetic gain by recurrent selection of important traits over many generations. However, the biological and genetic processes that allow continued genetic gain within a finite genepool are still unclear. For example, the Illinois maize long-term selection experiment achieved continuous increases in seed oil and protein concentration in a closed population for more than 100 generations without apparent loss of genetic variation (Dudley, 2007). Long-term trends in wheat (*Triticum aestivum* L.) breeding also reflect this, where significant genetic gain in traits such as yield has been achieved in the last century (Mackay et al., 2011; McCraig et al., 1995; Tadesse et al., 2019), whilst molecular studies have not found the expected reductions in genetic diversity over the same period of modern plant breeding (Fu, 2015; van de Wouw et al., 2010; White et al., 2008).

Selection on one trait can have positive or negative pleiotropic effects on other traits. For example, the Illinois long-term selection experiment found correlated responses to selection for oil and protein content and indirect effects on other traits, such as starch content. Wheat breeding requires selection for multiple traits of economic importance, including grain yield and quality traits, as well as other agronomically important or physiologically adaptive traits, such as developmental stage, plant architecture and disease resistance. In many cases, positive correlated responses in combinations of desirable traits can be achieved, but there are often complex trade-offs between antagonistically related traits. Considerable work has succeeded in identifying underlying quantitative trait loci (QTL) controlling individual yield components, such as grain size (e.g. Brinton et al., 2017) and spikelet number (e.g. Kuzay et al., 2019; Muqaddasi et al., 2019). However, the effects of such yield component loci rarely have consistent positive effects on yield in broader genetic backgrounds due to compensatory effects which trade-off against other yield components. For example, increased grain number per inflorescence in wheat is commonly associated with reductions in other yield components such as thousand grain weight, or tiller number (Corsi et al., 2021; Quintero et al., 2018; Xie and Sparkes, 2021).

In general, long-term increases in wheat yields have been achieved phenotypically by optimisation of harvest index (the ratio of grain to total shoot dry matter) to reduce intra-crop competition (Fischer and Kertesz, 1976), as well as through increased grain filling with starch carbohydrates (Lovegrove et al., 2020; Shewry et al., 2020). However, these have led to negative trade-offs in other valuable traits. Decreased competitive ability of modern wheat varieties with weeds (Murphy et al., 2008; Vandeleur and Gill, 2004) necessitates increased reliance on herbicides as well as potentially poorer uptake of soil nutrients (Ruisi et al., 2015). Yield loss from weed competition has become even more problematic in intensified cropping systems (Storkey et al., 2021). Additionally, increased yield and starch grain filling has been subject to the long-standing trade-off between yield and grain protein content (Simmonds, 1995; White et al., 2021), and has led to dilution of wheat grain protein content (Austin et al., 1980; Fufa et al., 2005) and mineral nutrient density (Davis, 2009; Shrewry et al., 2016). This has also led to higher optimum nitrogen fertiliser application rates to meet milling wheat grain protein requirements with diminishing increases in yield, and thus poorer nitrogen use efficiency (Hawkesford, 2014). Trade-offs between grain yield and both protein content and weed competitive ability seem not to have been generally addressed by commercial breeding due to yield being considered the highest economically important trait. Recent analysis by Raherison et al. (2020) suggested that negative pleiotropic genetic effects in wheat have rarely been compensated for and optimised by breeding, and Yang et al. (2022) showed that breeders’ selections have almost always been in favour of yield at the expense of protein. Changing economic, legislational, environmental and societal factors mean that breeding focus will increasingly need to consider how to deliver sustainable intensification of food supply, ensuring yield stability of our future crops in the face of such pressures. Plant breeding will play a role in delivering these goals, and will likely require the application of new breeding approaches and methodologies.

Genomic selection models aim to predict as large a proportion of heritable phenotypic variation as possible using genome-wide marker data to allocate estimated breeding values to untested individuals (Jannink et al., 2010; Meuwissen et al., 2001), and are likely to be a major source of improvement in plant breeding in the coming decades (Mackay et al., 2021). Genomic prediction models include genetic effects that don’t necessarily reach genome-wide significance in QTL mapping, which only detects large additive genetic effects and often fails to account for a large proportion of heritable trait variation in traits with complex genetic architectures, despite extensive genomic and phenotypic characterisation (Goddard et al., 2016). However, the role of non-additive epistatic effects in complex trait genetic architectures (i.e the interactions between genes) remains understudied, and is often overlooked (Carlborg and Haley, 2004) – likely due to the high computational requirements to model high order interactions (Jiang and Reif, 2015). Genomic prediction models that take epistatic effects into account have recently been developed, including the extension of the genomic best linear unbiased prediction (GBLUP) (Jiang and Reif, 2015) and machine/ensemble learning methods such as Random Forest (Schmalohr et al., 2018; Wright et al., 2016), which are often able to improve prediction accuracies in real datasets (Charmet et al., 2020).

The NIAB Diverse MAGIC (Multi-parent Advanced Generation Inter-Cross) wheat population (NDM) was recently developed to investigate the genetic architecture of a range of traits in wheat (Scott et al., 2021). It consists of 16 founders genotyped via exon and promotor capture sequencing and 504 recombinant inbred lines genotyped via whole-genome sequencing and imputation, resulting in ~1.1M single nucleotide polymorphisms (SNPs) between genotypes, or 55k SNPs after filtering for linkage disequilibrium (LD) (Scott et al., 2021). The 16 founders are wheat varieties that span 70 years of breeding and capture a large proportion of the northwest European genetic diversity. The genetic diversity present in the NDM is efficiently recombined through multiple generations of inter-crossing, eroding LD accumulated in the founders over long-term selective breeding. For this reason, traditional genomic prediction models, such as GBLUP, that make use of kinship relationships (Clark et al., 2011) may perform poorly in MAGIC where causal variants can be considered more independently (Scott et al., 2021). The NDM is ideal for investigating trait relationships, due to intensive phenotyping and the lack of the confounding effects of age and origin that are present in panels of selectively bred varieties (Scott et al., 2020). Furthermore, this population provides a good test for multi-trait selection indices, such as grain yield protein deviation (GYPD; Michel et al., 2019), that have been proposed to help minimise trade-offs between traits.

Here we use NDM resources as a microcosm of long-term selection in wider wheat breeding programmes to test differing approaches to selection and genetic models. Within the overall context of understanding the phenotypic and genetic mechanisms that may enable enhanced genetic gain in the future, we (i) investigate complex trait relationships relating to yield in the observed population of lines. We then (ii) develop multi-trait genomic prediction models that increase prediction accuracy by exploiting pleiotropic effects among traits, and (iii) investigate how increased prediction accuracy translates to greater genetic gain in simulation of long term-recurrent genomic selection within the NDM. We also (iv) simulated both phenotypic and genotypic effects of recurrent phenotypic selection within the population comparing different true genetic models based on genomic prediction models trained on the observed population. Our results reveal correlated responses in a wide range of traits when selection is purely on yield, as well as the potential to achieve genetic gain in several traits of interest that trade-off by using use multi-trait selection indices. Comparison of response to selection under differing genomic prediction models (simplified models with a minimal number of additive effects versus more complex polygenic models that take higher order epistatic interaction effects into account) also highlights the important role of both pleiotropy and epistasis as potential mechanisms for continued genetic gain in crop breeding.

## 2. Methods

### 2.1 Germplasm, phenotypic and genotypic datasets

Genotypic and phenotypic data for the NDM wheat population was sourced from Scott et al. (2021). Briefly, the population of 504 NDM recombinant inbred lines derived from the 16 founders was phenotyped for a wide range of traits over two successive seasons (2016-2017 and 2017-2018) in the United Kingdom (UK). All but one of the 73 traits described by Scott et al. (2021) (**Table 1**) were used, the exception being seed germination rate (GR) due to a large proportion of missing data. Traits measured in each year were considered separately. Missing data for all remaining traits (at <1.2%) were imputed with the median trait value. Line genotypes were previously characterised by skim sequencing and imputed using the founder haplotypes (Scott et al., 2021). Of 1.1M SNPs identified from founder exome and promoter sequencing, we use the subset of ~55k SNPs after pruning for LD for our analyses. Missing marker data (~1%) were imputed using the ‘missForest’ package (Buhlman, 2011) in R (R Core Team, 2020), which uses non-parametric Random Forest prediction models to iteratively predict and impute missing data on a marker-by-marker basis.

**Table 1.** Abbreviations of traits phenotyped in the NDM, as described by Scott et al. (2021). All data are from field trials, except where noted. Nursery = data collected from 1×1m unreplicated plots. Field = data collected from 2×6m replicated plots. Trait groups indicate groups of strongly positively or negatively correlated traits that grouped together through hierarchical clustering as shown in **Figure 1**. Some traits were phenotyped at multiple time points (GLA) and in both trail years so that a total of 72 traits were included.

| Abbreviation | Trait | Trait group | Abbreviation | Trait | Trait group |
| --- | --- | --- | --- | --- | --- |
| PHS | Pre-harvest sprouting | 1 | GW | Grain width | 6 |
| PIG | General pigmentation | 1 | HET | Height to ear tip | 6 |
| SW | Specific weight | 1 | HFLB | Height to flag leaf base | 6 |
| FLS | Flag leaf senescence | 2 | LOD | Lodging | 6 |
| GS39 | Flag leaf emergence date | 2 | TGW | Thousand grain weight | 6 |
| GS55 | Ear emergence date | 2 | TIS | Tip infertile spikelets | 6 |
| GS65 | Anthesis date | 2 | AWN | Presence of awns | 7 |
| JGH | Juvenile growth habit (Nursery) | 2 | EL | Ear length | 7 |
| SH | Spring habit | 2 | ETA | Ear taper | 7 |
| FLA | Flag leaf angle | 3 | GLAU | Glaucosity | 7 |
| FLF | Flag leaf floppiness | 3 | SPIG | Stem pigmentation | 7 |
| FLL | Flag leaf length | 3 | BIS | Basal infertile spikelets | 8 |
| GLA# | Green leaf area (10 time points) | 4 | EW | Ear weight | 8 |
| GPC | Grain protein content | 5 | FLW | Flag leaf width | 8 |
| GY | Yield | 5 | GPE | Grains per ear | 8 |
| FLED | Flag leaf to ear distance | 6 | GPS | Grains per spikelet | 8 |
| GA | Grain area | 6 | TS | Total spikelets | 8 |
| GL | Grain length | 6 | YR | Yellow rust infection (Field and Nursery) | 8 |

### 2.2 Statistical analysis

All analyses were conducted using R statistical analysis software. Pearson’s correlation coefficients among all investigated traits were calculated. Hierarchical clustering of traits was performed using the ‘hclust’ R function and ‘complete’ method, where the distance matrix (*d*) was derived from the equation:

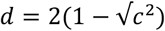

where *c* represents the trait correlation matrix. Traits were then assigned to eight groups using the ‘cutree’ R function.

#### 2.2.1 Genomic prediction models

Two contrasting genomic prediction models were compared for both single-trait (ST) and multi-trait (MT) models. These included generalised linear models including the Lasso penalty (LASSO) implemented in the ‘glmnet’ R package (Friedman et al., 2010) where the majority of SNP effects are shrunk to zero. For comparison, a non-linear, statistical learning approach was also used which generally included much larger numbers of SNPs with non-additive effects in each model: Random Forest (RF), implemented in the ‘randomForest’ R package (Breiman, 2001). For LASSO models, the value of lambda (a shrinkage penalty) used for each prediction was optimised using 8-fold cross validation. For RF models, 300 trees were run per model and default parameters of one third of variables randomly sampled at each split, and a minimum of five observations in terminal leaf nodes were used. Previous work by Scott et al. (2021) found that ridge regression models that include all marker effects with a small, but non-zero effect, did not have as high prediction accuracy as LASSO in the MAGIC population, and so were not further tested here.

Two types of MT models were implemented. Firstly, by performing single value decomposition (SVD) of the matrix of all phenotypes, as proposed by Montesinos-López et al. (2019a), whereby each of the decomposed and uncorrelated vectors from all the traits were predicted as traits themselves using the same genomic models as for ST predictions. The predictions of vectors were then back-transformed to the original trait scales to derive the MT predictions per-trait. Secondly, a multi-trait stacked ensemble method was also used which employs an approach often used in machine learning (Spyromitros-Xioufis et al., 2016) and has previously been applied for Bayesian multi-output regression of multi-trait predictions (Montesinos-López et al., 2019b; Sapkota et al., 2020). For this, a two-step model was used where each trait was first predicted from genomic data with the same genomic models as for ST predictions, and then all trait predictions were used as explanatory variables (features) in a second multi-trait ensemble model to again predict each response trait. Both first and second stage predictions were fitted only on data from the training fraction and predictions were independently made for test lines with only genetic marker data. Either LASSO or RF models were used to fit first stage ST models, but only RF models were used to flexibly include non-linear multi-trait relationships for the second stage ensemble models. Information from related traits is therefore used in a model, trained only on the training fraction to adjust single-trait predictions made directly from genomic data. Unlike trait-assisted genomic prediction, such as used by Fernandes et al. (2018), no observed trait data are used in the tested cross-validation fraction. As both MT prediction approaches can be applied with any genomic prediction model for each ST prediction or for SVD vectors, we were able to compare LASSO and RF genomic models for both ST and MT approaches.

Prediction accuracies were determined by performing three rounds of 10-fold random cross-validation among all lines in the dataset and averaging the three Pearson’s correlation coefficients between observed and predicted trait values across all cross-validation folds. Valid comparisons of prediction accuracy were ensured by testing all prediction models using the same cross-validation fold assignments. After model cross-validation, full prediction models were fitted using the entire dataset for combinations of both ST and MT models with both LASSO and RF genomic models. Variable importance scores for each SNP marker in RF genomic prediction models and for each trait covariate in MT ensemble models were derived from the full models as the mean decrease in mean square error using the ‘importance’ function in the ‘randomForest’ R package. Effect sizes for each SNP marker were also derived from full LASSO models where the majority of SNP effects were shrunk to zero.

#### 2.2.2 Simulations of recurrent genomic selection

We first simulated a recurrent genomic selection programme within the NDM to assess the performance of different prediction models to achieve long-term genetic gain in grain yield. This was done within a framework of assuming a true inherited genetic model based on the MT ensemble RF genetic model as outlined above and trained on the observed genetic and phenotype data. True phenotypes were derived from predictions from this model for genotypes at each cycle of simulations and the different genomic prediction models outlined above were trained on the individuals in the first cycle. New cycles of genotypes derived from crossing selected fractions of lines were simulated using a genetic map of ~55,000 SNPs.

The genetic map (Supplementary Table S1 and Supplementary Figure S1) was constructed using the ‘qtl2’ R package (Broman et al., 2019) with the marker data ordered by physical map position (RefSeq v1,0, IWGSC, 2018). The genetic map distance was then re-estimated using the ‘est_map’ function with 1,000 maximum iterations and an assumed genotyping error probability of 0.001. The cross object was considered as a 16-way multi-parent recombinant inbred line population, so the differing local recombination effects for each founder haplotypes were preserved for subsequent simulations. 23 markers were removed from the full set which caused genetic map distortion.

Selection of lines at each generation were made based on predicted phenotypes from the genomic prediction model. To reduce excessive inbreeding and loss of genetic variance, 15 lines from different 16-way or bi-parental families with the highest selection index values were selected. 30 offspring inbred line genotypes were then simulated for each of 105 possible pairwise cross combinations among the selected lines so that the following generation comprised of 3,150 lines from 105 biparental families. The phenotypes of these were again predicted from the genomic prediction models trained on the true phenotypes of the first generation and the process repeated for 20 cycles of recurrent selection. 20 iteration repeats of the simulations were run for each genomic prediction model. Genomic prediction models were fitted as detailed above for ST and MT, LASSO and RF models, but additionally RF models were run that were restricted to a tree depth of one (RF1) to completely limit the degree of marker interaction effects. 2,000 trees were used for RF1 models.

Genetic gain for each trait over the selection simulations were determined by comparing the mean true trait value of all lines at each generation to the mean true trait value in the first generation. The divergence from this mean among different traits was standardised to the standard deviation of trait values in the first generation.

The accuracy of genomic prediction models was also determined as the Pearson’s correlation coefficient between the true and predicted trait values among genotypes at each simulation cycle.

#### 2.2.3 Simulations of recurrent phenotypic selection

In addition to simulations of different genomic selection procedures within a simulated true genetic model, we also compared simulations with different true genetic models to assess both phenotypic and genomic response to selection with different genetic model assumptions of trait genetic architecture. Simulations were run as above but selections of individuals were based on the true phenotypes derived from different genetic models so that it was assumed that the simulated breeder could make perfect estimates of trait values from phenotypic selection. Different simulations were run for ST and MT as well as RF and LASSO models as outlined above and for different selection indices as outlined below. Genetic response to selection was also characterised as the change in allele frequency for all ~55,000 SNPs at each generation, and the genetic diversity was calculated as the number of polymorphic SNPs at each generation. For each set of simulations, traits or SNP markers were considered under selection rather than drift if their response to selection was significantly different to 0 considering all 20 simulation repeats using a t-test.

#### 2.2.4 Selection indices

Indices for simulated selection were defined as follows:

1. Grain yield measured in each trial year.
2. Multi-trait index including grain yield and traits known to be associated with weed competitiveness. The weed competitive ability selection index (Weeds_ESIM) was based on the restricted eigenvector selection index method (RESIM) (Cerón-Rojas et al., 2008). For this, principal component analysis was performed on a selection of desirable traits based on the literature, which included yield measured in both years as well as traits previously identified as valuable for weed competitive ability. These included high early vigour, measured as green leaf area (GLA) over the development phase before flowering time, and planophile (horizontal) flag leaf angle (FLA) (Andrew et al., 2015; Kissing Kucek et al., 2021; Korres and Froud-Williams, 2002; Mwendwa et al., 2020). To mitigate risk of lodging (i.e. the permanent displacement of a stem from vertical), mean crop height (HET) between both years was also then restricted to values between 60-65 cm. The vector weightings on the first principal component with mean HET values between this range therefore represented the Weeds_ESIM. Most traits involved in this selection index were positively correlated so the first principal component could be assumed to provide a desirable combined index for selection.
3. Grain yield protein deviation (GYPD). The GYPD selection index was calculated as the sum of the scaled and centred mean yield and protein across both years using the equation:

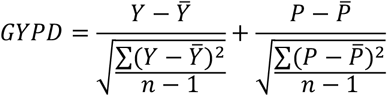

where *Y* and *P* is the mean line yield and protein across years, respectively.

## 3. Results

### 3.1 Grain yield is correlated with multiple traits in the observed population

We analysed data from a genetically diverse and highly recombined 16-founder wheat MAGIC population for a wide range of agronomically important traits over multiple trial years (72 trait – year combinations) to investigate complex trait-trait relationships and their implications for breeding. Correlation analysis across all traits and years revealed complex trait relationships and substantial differences in patterns between the two trial years **(Figure 1** and Supplementary Table S2). Considering grain yield (GY) as the primary trait of interest, many other secondary traits were found to be correlated (**Figure 1**). The strong negative trade-off between yield and grain protein content (GPC) was mediated by yield component traits. For example, grain size traits (such as thousand grain weight; TGW), grains per spikelet (GPS) and total spikelets per ear (TS), were all positively correlated with GY, but negatively correlated with protein content and with each other. Therefore, potential benefits of selecting for any one of the yield component traits in isolation are buffered by problematic trade-offs with other yield component traits and likely have negative effects on protein content.

**Figure 1.**
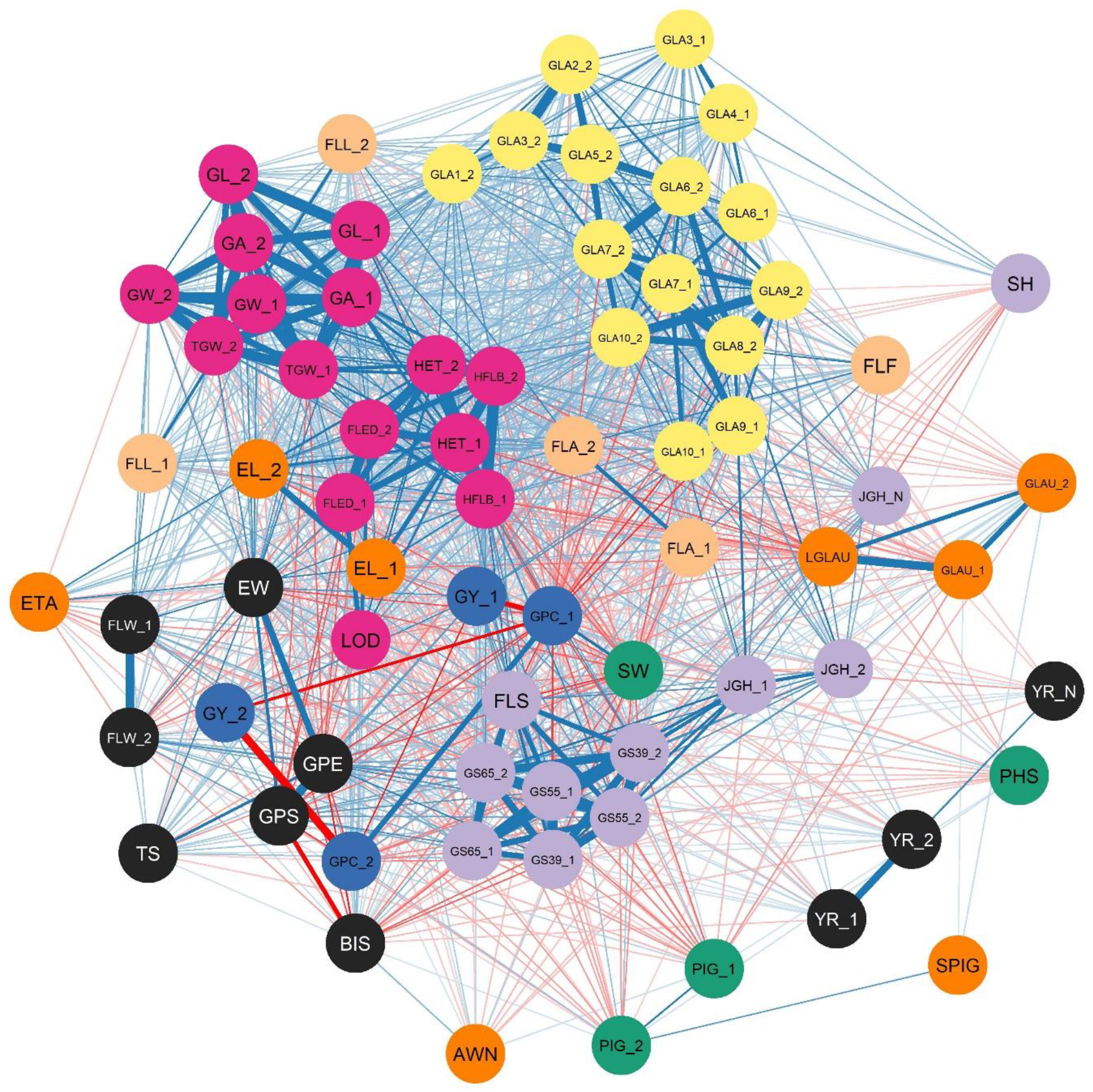
Correlation network for 72 traits measured in two trial years among 504 NIAB Diverse MAGIC lines. Grain yield = GY. Abbreviations for all additional trait names are given in Table 1. Trait node colours indicate the eight groups of related traits as identified using hierarchical clustering. The _1 and _2 designations used after trait abbreviations refer to trial year 1 and trial year 2, respectively. Blue and red connecting lines indicate positive and negative correlations, respectively, while line thickness is relative to correlation *p*-value significance.

Differential relationships between yield and other developmental stage and plant architecture traits between the two trial years were also found. In year 1, taller and later flowering genotypes were generally higher yielding (yield – height to ear tip correlation = 0.20; yield – heading date correlation = 0.32), whereas in year 2, the correlation between height and yield was negative (correlation = −0.11) and between yield and heading date was non-significant. Therefore, strong genotype-by-environment interaction (G×E) effects on yield means that selection for yield, or related adaptive traits, in any single environment may have limited potential to increase yield in another environment. However, contrasting patterns of rainfall and temperature between the two yield trial years (**Figure 2**Figure 5), in which year 1 was characterised by high temperatures and drought before anthesis (March and April) whilst year 2 was characterised by extreme terminal heat and drought after anthesis (June and July), may explain the differences in relationships between adaptive traits and yield in this study. Both of the two trial years experienced different extremes of monthly climate variables compared to the distributions of the last 56 years, so were considered separately in these analyses.

**Figure 2.**
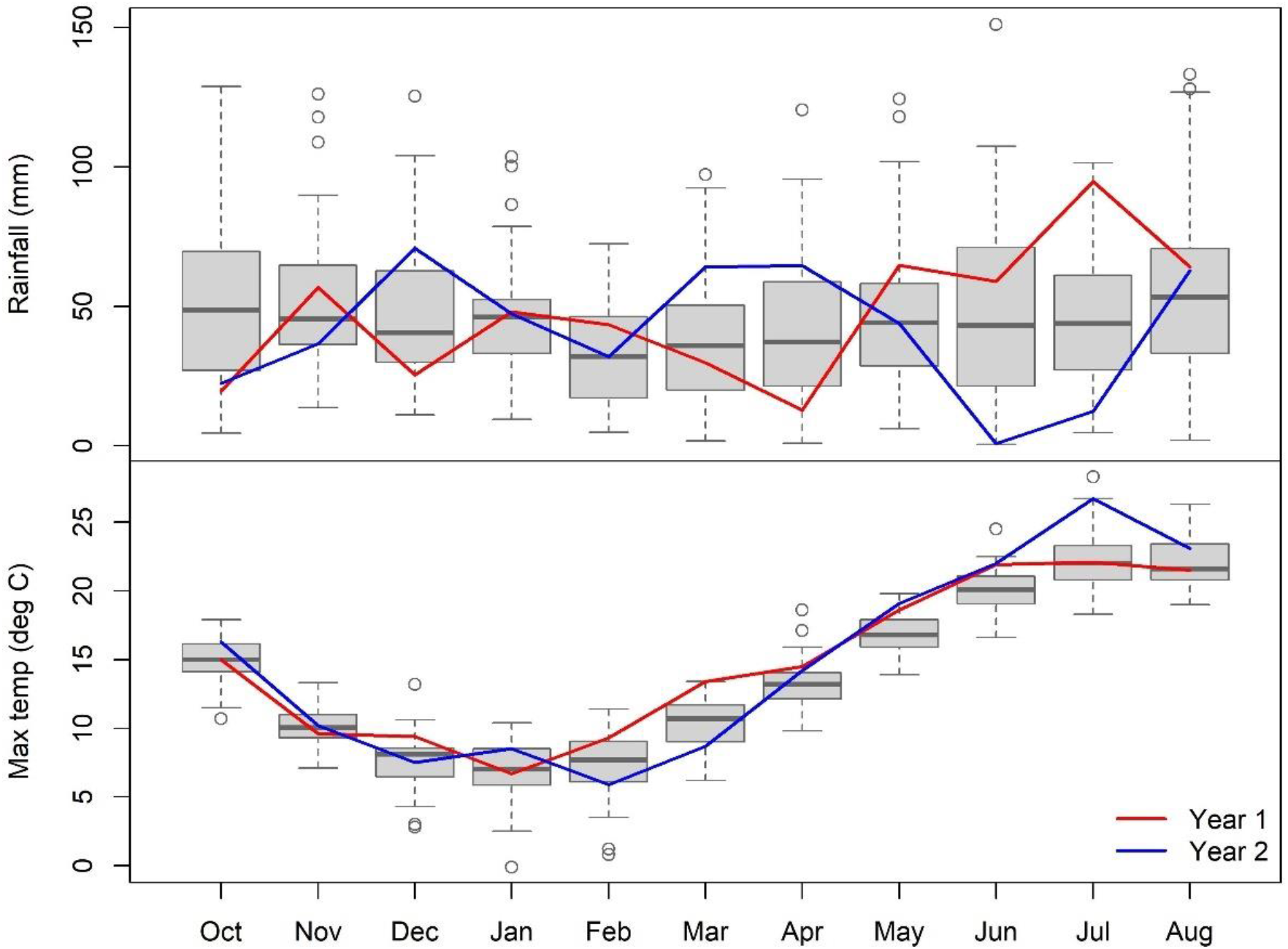
Contrasting patterns of monthly rainfall and maximum temperature over the growing season for the two trial years. Boxplots indicate historic variation in data for each month from 1960 to 2016 (horizontal line = median, boxes = interquartile range, whiskers = 1.5 times the interquartile range and points = values outside 1.5 times the interquartile range).

Other plant architecture traits, such as green leaf area (GLA) in the development phase, juvenile growth habit and flag leaf morphology had weak but positive correlations with yield, suggesting potential relevance of these traits as mechanisms to increase yield, or as valuable traits themselves to select for in combination with yield to increase crop competitive ability with weeds. However, the strong positive correlations between GLA traits and plant height traits mean that increasing these traits without increasing lodging risk may be problematic. Optimising combinations of important traits therefore requires consideration of correlated responses due to pleiotropy and linkage.

### 3.2 Genomic prediction of complex traits

We tested the accuracy of several genomic prediction approaches to determine the genetic architecture of the multiple related traits, using both single-trait (ST) and multi-trait (MT) models that take into account relationships among correlated traits. LASSO represents models including trait genetic architecture controlled by a minimal number of additive genetic effects across the genome, while Random Forest (RF) represents models including a much greater number of interacting genetic effects. RF outperformed LASSO for most traits in ST models and was particularly advantageous for traits with generally low genomic prediction accuracy, such as GLA and grain yield in both years (**Figure 3a**). Prediction accuracy was increased from 0.34 and 0.20 in ST LASSO models to 0.38 and 0.27 in ST RF models for grain yield measured in each year respectively. In contrast, traits with greater prediction accuracy, such as plant height or grain dimension traits, were better predicted by LASSO. This suggests that RF can successfully predict genetic effects in traits with more complex genetic architecture, potentially using non-additive and epistatic effects.

**Figure 3.**
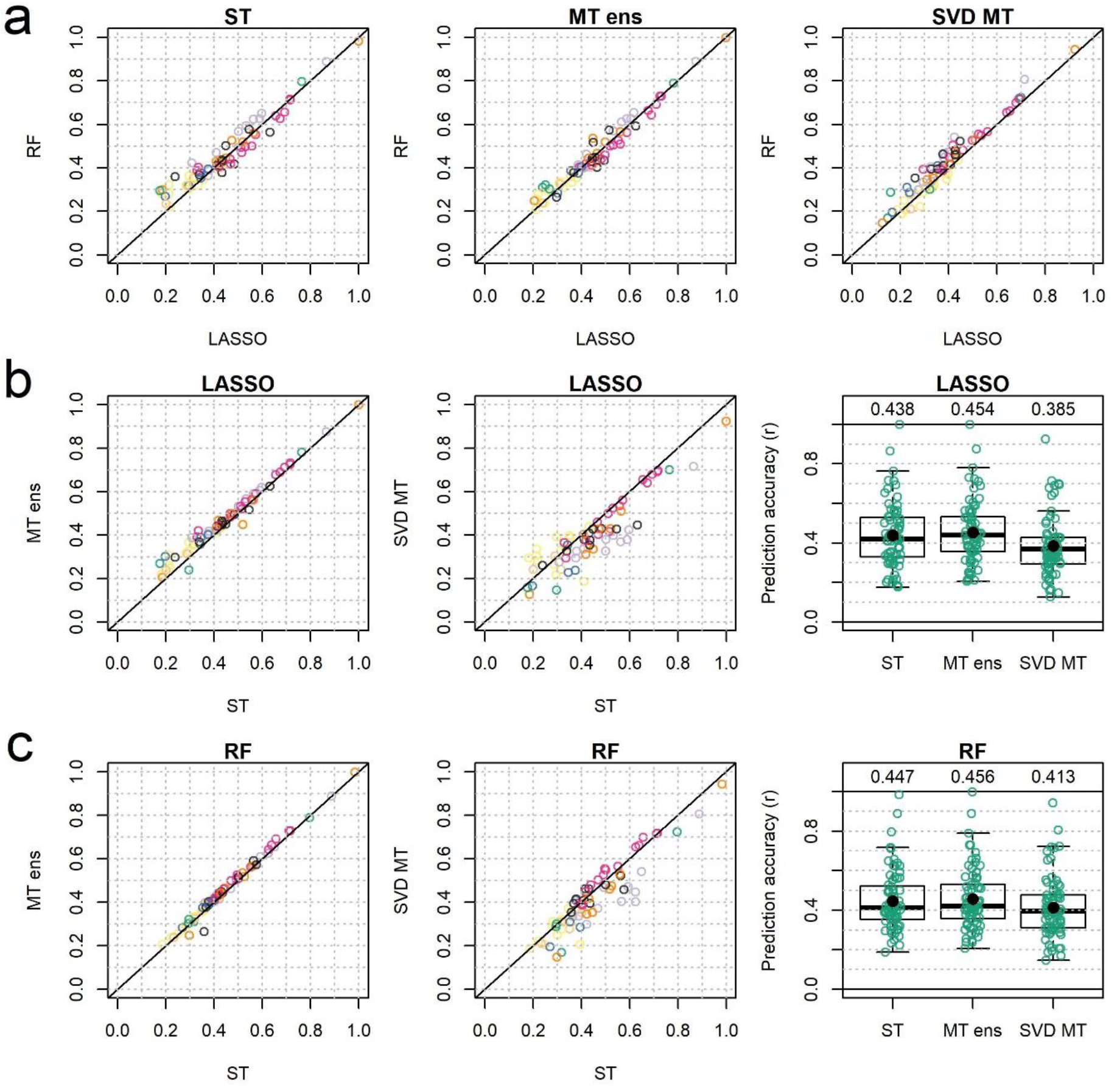
Comparison of genomic prediction accuracies for all traits. Each circle represents a trait-year combination, with circles colour coded according to trait group, as in **Figure 1**) between LASSO and Random Forest (RF) models using single-trait (ST), multi-trait ensemble (MT ens) and Single Vector Decomposition (SVD) approaches. Row (a) compares LASSO and RF prediction models, row (b) compares ST with both MT approaches for LASSO prediction models, and row (c) compares ST with both MT approaches for RF prediction models. Horizontal lines in boxplots represent the median and black dots represent the mean prediction accuracy across all traits, which is also shown above each boxplot.

Multi-trait ensemble (MT ens) models consistently produced more accurate predictions for almost all traits for both RF (86% of traits) and particularly for LASSO (90% of traits) models (**Figure 3b-c**). On the other hand, single vector decomposition (SVD) MT models were less reliable and generally resulted in lower prediction accuracies (**Figure 3b-c)**, although RF notably outperformed LASSO in SVD MT models. Therefore, including information from predictions of other related traits in a multi-trait ensemble model was advantageous over attempting to predict pleiotropic effects directly in a decomposed trait matrix. Traits that were poorly predicted by LASSO compared to RF for ST models (**Figure 3a**, left panel) had particularly increased prediction accuracies via the MT ensemble predictions for LASSO (**Figure 3b**, left panel). This suggests traits that were predicted better by either of the multi-trait models have few direct or large genetic effects and are rather the culmination of many other component traits. These indirect genetic effects may be picked up in complex RF prediction models, but are best captured by the use of MT models that can directly model pleiotropic trait trade-offs

MT ensemble models increased genomic prediction accuracy of grain yield from an average of 0.27 to 0.33 for LASSO models and from 0.32 to 0.34 for RF models on average across both years and cross-validations. Variable importance of trait covariates in the Random Forests used in the MT ensemble models indicates the influence of these traits in the model. Across all traits, most highly influential traits had strong positive or negative correlations in each year among the observed lines (**Figure 4a**). Considering grain yield in each year as the primary trait of interest, highly correlated traits such as GPC and grain yield measured in the other year, were highly important in MT ensemble models for grain yield, suggesting that pleiotropic effects mediating the grain yield and protein content trade-off are useful for predicting grain yield itself (**Figure 4b**). Base model predictions of yield in the other year as the focal yield trait were also included in models with high importance suggesting that the ensemble model effectively takes yield G×E effects into account. Developmental stage traits including dates of growth stages GS39 (flag leaf blade all visible), GS55 (ear half emerged) and GS65 (flowering half-way complete) and flag leaf senescence (FLS) were particularly important covariate traits and were positively correlated with yield in year 1 to a greater extent than in year 2 (**Figure 1**). These traits also featured with greater importance in MT ensemble models when grain yield in year 1, compared to year 2, was predicted **(Figure 4b**). This indicates that later-developing lines were predicted to be higher yielding in the year without terminal drought stress. Other yield component traits including grains per spikelet (GPS) and grains per ear (GPE), but not grain size traits, were found to be of high importance (**Figure 4b**). Many GLA traits, particularly when measured in the spring, were also included with fairly high importance in MT ensemble models (**Figure 4b**), suggesting a role of the crop development phase in resource acquisition for final grain yield. These results not only identify important traits for inclusion in multi-trait prediction models, but also physiological mechanisms for grain yield improvement.

**Figure 4.**
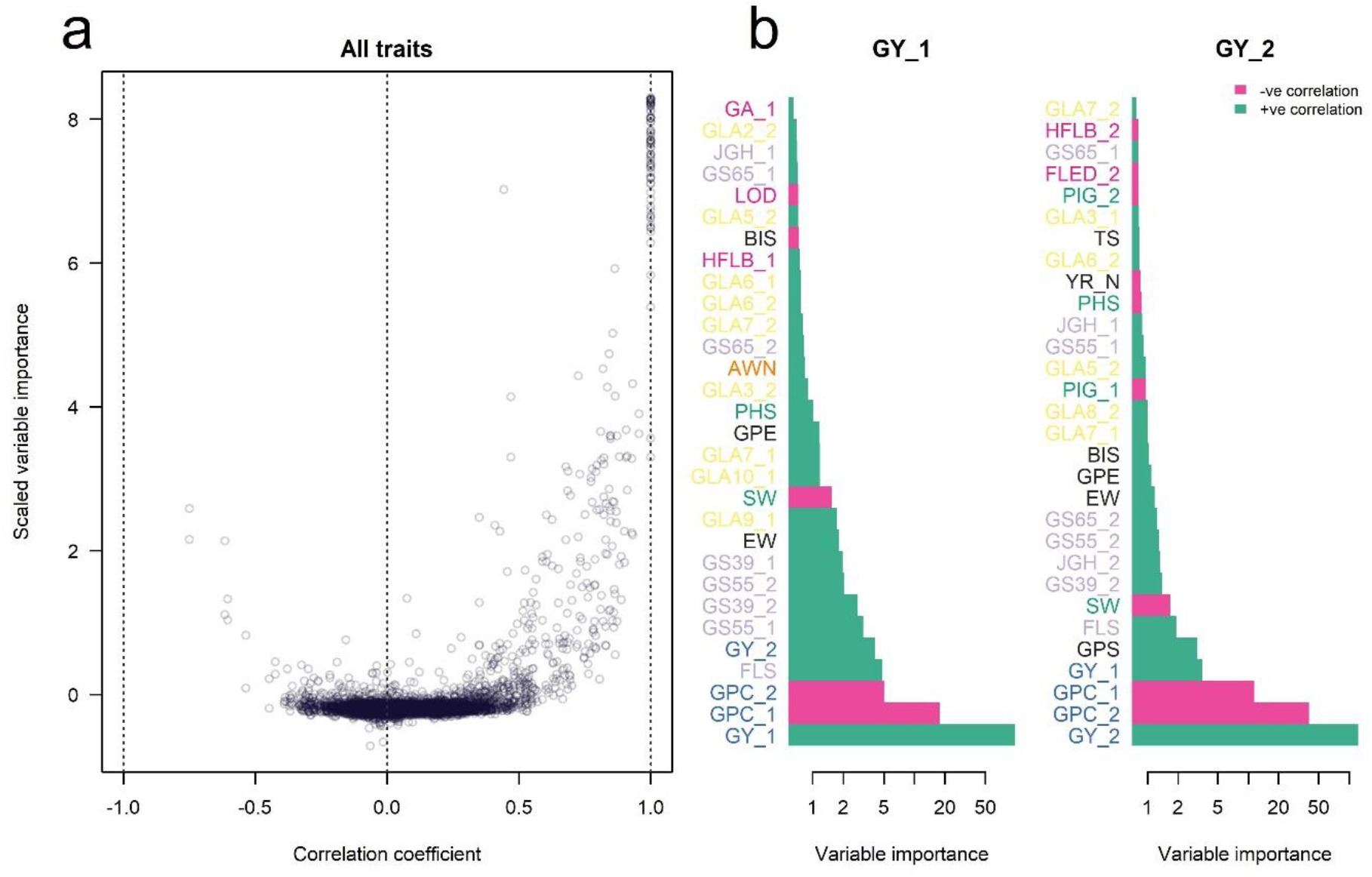
The Influence of related traits in multi-trait ensemble prediction models. (**a**) The relationship between pairwise correlation coefficients among all traits and years and the variable importance score in Random Forest multi-trait ensemble models for all target traits. (**b**) The 30 most important trait variables used in Random Forest multi-trait ensemble models for prediction of grain yield in each year (GY_1 and GY_2). All trait abbreviations are as listed in **Table 1** and colour coded according to trait group, as in **Figure 1**. The _1 and _2 designations used after trait abbreviations refer to trial year 1 and trial year 2, respectively.

### 3.3 Accurate genomic prediction models increase long-term genetic gain in simulation of recurrent genomic selection

We then investigated the potential for different genomic prediction models to achieve genetic gain in yield through simulation of a recurrent genomic selection programme within the NDM. MT ensemble RF models were found to be the most accurate model from cross validation within the observed population (**Figure 3**) and so were used as the true genetic model to define the true phenotypes of simulated lines. The genomic prediction models were then trained on the simulated true phenotype data of lines in the first generation and genomic predictions of phenotypes were used to make selections for subsequent cycles of lines.

The accuracy of genomic prediction models over the course of the selection simulations generally reflected those in cross validated of the observed data. Prediction accuracy of all models decreased in later cycles of the simulations, but models that were more accurate in the observed data and maintained accuracy for longer, such as MT RF and ST RF (**Figure 5a**), achieved greater long-term genetic gain in yield (**Figure 5b**). RF models that included restricted trees with an interaction depth of one so that genetic marker interaction effects could not be included were much less accurate and led to less genetic gain, particularly for grain yield in year 1 (**Figure 5**). This suggests an important role for prediction of non-additive genetic effects in RF models for continued accuracy of genomic prediction models, particularly as breeding cycles become more distantly related to the training set.

**Figure 5.**
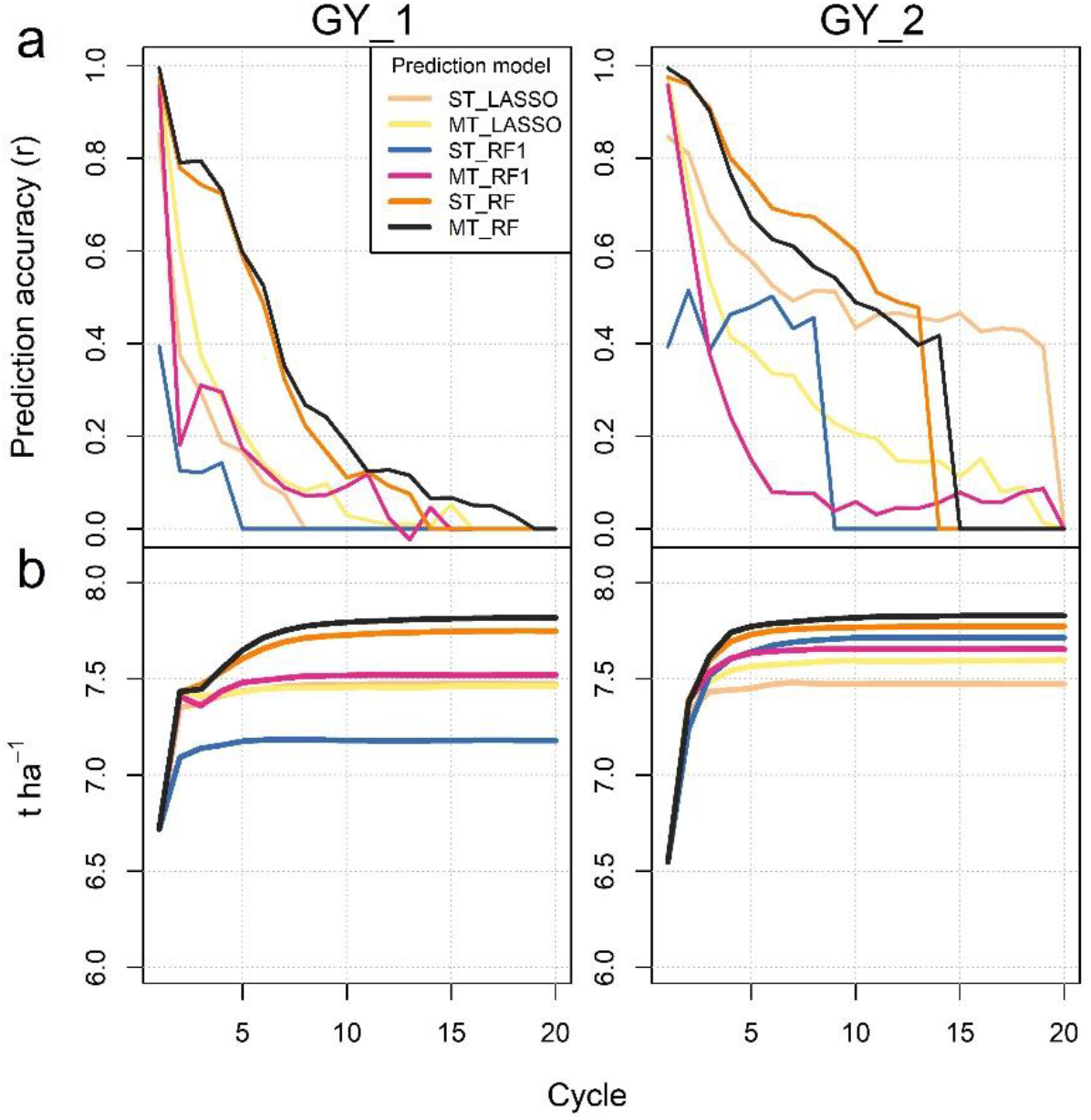
Trends in accuracy of (a) genomic prediction models and (b) resulting genetic gain over a simulated recurrent genomic selection programme for grain yield (GY) measured in two years trial years. Genomic prediction models include single-(ST) and multi-trait (MT) ensembles models for LASSO and Random Forest (RF). RF1 indicates Random Forest models with a restricted interaction depth of one. Lines represent the averages across 20 simulation repeats.

### 3.4 Simulation of recurrent phenotypic selection for grain yield reveals indirect effects on multiple traits

Phenotypic correlations identified traits that may be under similar genetic control through either pleiotropy or linkage, so we investigated the potential for recurrent selection to achieve genetic gains in traits directly under selection as well as indirect effects on other traits. Simulations of a phenotypic recurrent selection programme were run that compared different selection indices and true genetic models. For these, selections were made based on the true phenotypic values rather than the predicted phenotypes as for simulation of recurrent genomic selection.

Firstly, we simulated selection based purely on grain yield measured in each trial year. Considering the MT ensemble RF as the true genetic model, which achieved the greatest prediction accuracy across traits (**Figure 3**),selection on grain yield *per se* resulted in rapid genetic gain in the yield trait under direct selection as well as indirect effects on other related traits (**Figure 6**). These included selection for combinations of related traits that were complementary to, as well as those that were antagonistic to, their correlation in the unselected population. As an example of complementary trait selection, grain protein content (GPC) was shown to be strongly negatively correlated with grain yield in the original population (**Figure 1**) and therefore rapidly decreased as grain yield was selected for. Although both grain yield and GPC are both positively valued traits, here we define these as under complementary selection were their trait correlations and selection covariance are in the same direction. In contrast, antagonistic trait selection could be demonstrated by plant height traits and grain dimension traits that were all positively correlated with each other in the original population (**Figure 1**), but covaried negatively over time in the simulated population under selection for yield measured in each trial year; large grain size traits increased over time (GA, GL, GW, TGW), whereas plant height traits (HET, HFLB) decreased (**Figure 6**). Similarly, green leaf area (GLA) traits over the foundation development stage were generally positively correlated with plant height traits but increased over time as grain yield was selected for, while plant height traits decreased (**Figure 6**). Although the majority of trait relationships had complementary rather than antagonistic trait correlations in the original population and covariances in the simulated population (55.4 and 57.8% of pairwise relationship when grain yield selected in each year respectively had both positive or both negative correlations and covariances under selection; **Figure 6**), the significant remaining proportion did not. This indicates that antagonistic trait trade-offs were required to be optimised to achieve the genetic gains in yield simulated in the population and highlights the benefit of the multi-trait prediction approach.

**Figure 6.**
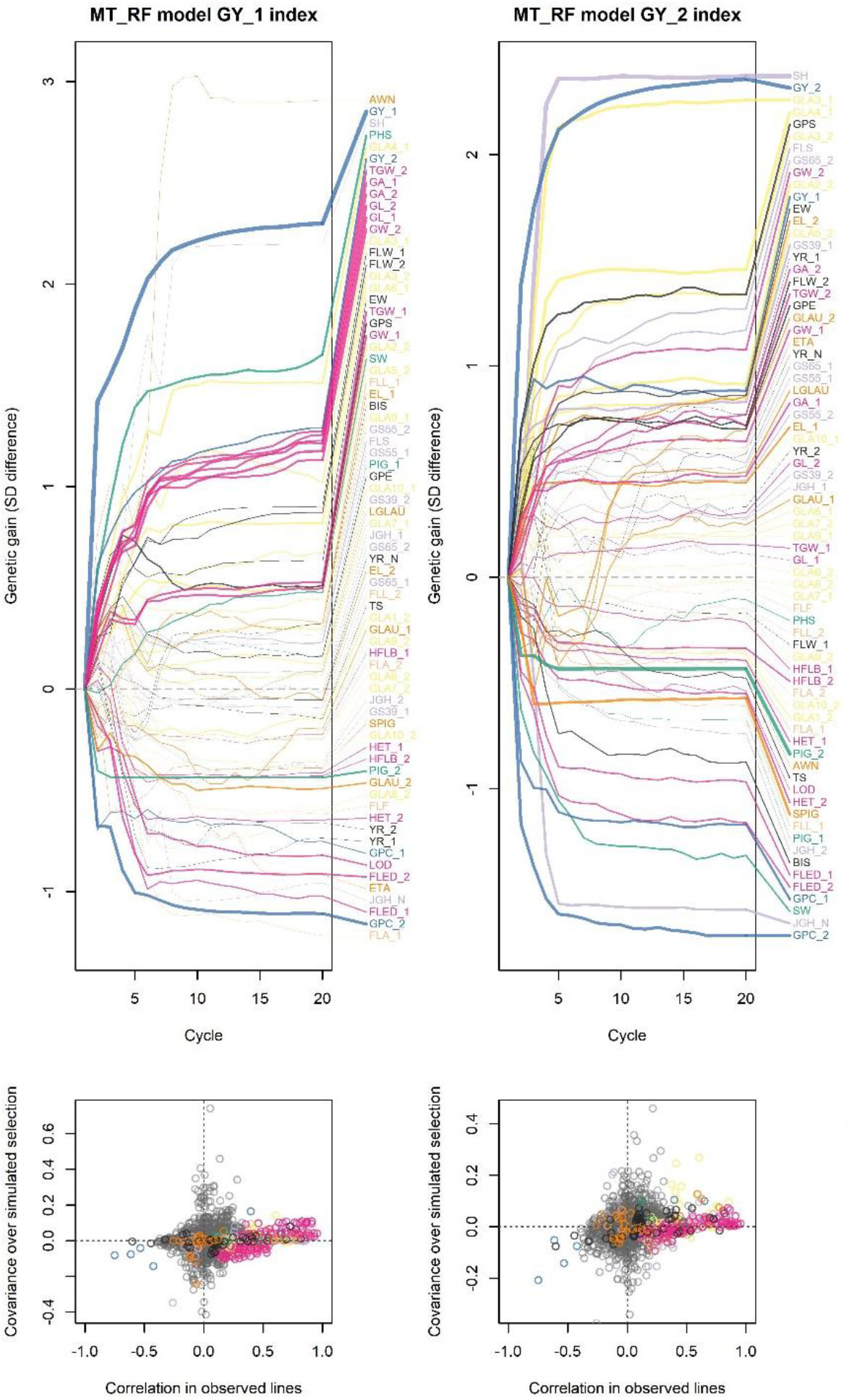
Simulated phenotypic response to selection on grain yield measured in two years. Upper plots indicate response of all 72 traits under simulated recurrent selection for grain yield in each of the two trial years based on multi-trait Random Forest (MT RF) genomic prediction models. Genetic gain was calculated as difference in population mean trait values to generation one and scaled to the standard deviation of the trait values in generation one. Line colours relate to trait groups identified by hierarchical clustering shown in **Figure 1**. The _1 and _2 designations used after trait abbreviations refer to trial year 1 and trial year 2, respectively. Line widths are relative to the t-test significance of each trait genetic gain from cycle 0 to 20 across all 20 simulation repeats. Lower plots compare the correlations between all pairs of traits in the original population and the covariance between trait pairs over time in the simulated population under selection. Points in upper right or lower left quadrants indicate both positive or negative correlation and covariance which demonstrates complementary trait selection. Points in the lower right or upper left quadrants represent differing positive or negative correlation and covariance, suggesting antagonistic trait selection. Point colours indicate pairs of traits that are both in the same trait group following the colour scheme in **Figure 1**, while grey points indicate trait pairs from different groups.

As suggested by the low correlation between yields in each year, selection for yield in either year had only limited effects on yield in the other year where approximately half the genetic gain in yield in the alternate year was achieved in simulated selection for yield in either year. This G×E effect for yield was reflected by how the yield component traits were co-selected with yield between the two years. Grains per ear (GPE) and grains per spikelet (GPS) were increased when selection was for grain yield in year 2 but remained mostly neutral for grain yield in year 1 (**Figure 6**). Further to the differential importance of traits in the multi-trait ensemble models outlined above, differential selection responses of yield component traits according to yield in differing environments highlights the capacity for G×E interactions to buffer response to selection for grain yield. However, when G×E is predictable, in certain target environments, contrasting yield component strategies could be used to adapt the crop to the environment. For example, it may be supposed that a genotype that can be high yielding by producing many grain sites per ear throughout an extended development phase due to being later flowering would be better adapted to environments without terminal drought stress.

### 3.5 Selection indices enable optimisation of trait trade-offs

We next tested whether multi-trait selection indices could be employed to simultaneously optimise selection for yield and other traits of interest, such as grain protein content (GPC) or crop architecture traits that aid competition with weeds. As outlined by the observed trait correlations, early season green leaf area (GLA) traits and grain yield were found to be slightly positively correlated, so could be co-selected, but the additional association between GLA and plant height would need to be restricted to limit risk of lodging. We therefore simulated effects of a phenotypic selection strategy based on selection index to increase important traits for crop competitive ability with weeds in combination with grain yield, whilst restricting changes in plant height as well as an index for high grain yield protein deviation to combine both negatively correlated traits.

Considering the ST RF as the true genetic model, the combined grain yield + weed competition selection index succeeded in increasing desirable competitive traits including GLA, flag leaf area (FLA) as well as grain yield in both years, whilst maintaining plant height at an acceptable level (**Figure 7**). Indirect effects on other traits included rapid early selection for spring-type growth habit (SH) up to the fifth breeding cycle, but which then remained at around 90% frequency in the population without fixation in any of the of the simulation repetitions. The GYPD selection index also achieved genetic gain in desirable traits (grain yield and protein content in both years) and had some indirect effects on related traits (**Figure 7**). As an example of one trait that was co-selected with GYPD, flag leaf width (FLW) increased in all simulation repeats, increasing by 3% and 4.2% for the trait when measured in year 1 and year 2, respectively. Whilst most of the trait relationships selected for in the weed competition index were positively correlated and complementary traits, such as all of the GLA traits and grain yield, the GYPD selection index included more antagonistic trait relationships (positive correlation and negative covariance or negative correlation and positive covariance) that were required to be optimised in addition to yield and protein trade off (**Figure 7**). For example, under GYPD selection, grains per spikelet (GPS) correlated negatively with GPC in each year (correlation = −0.33 and −0.36 in each year respectively), but covaried positively over simulated selection (covariance = 0.52 and 0.35 in each year respectively), where GPS increased by an average of 0.23 over the course of simulated selection while GPC measured in each year also increased by 2.48% and 1.57% respectively. Furthermore, the flag leaf to ear distance (FLED) and GPC measured in year 2 correlated positively (correlation = 0.21), but covaried negatively over the simulated selection (covariance = − 0.25), where FLED decreased by 2.73cm while GPC increased by 1.57% over the course of simulated selection. These provide examples of trait mechanisms by which yield and GPC could be simultaneously selected.

**Figure 7.**
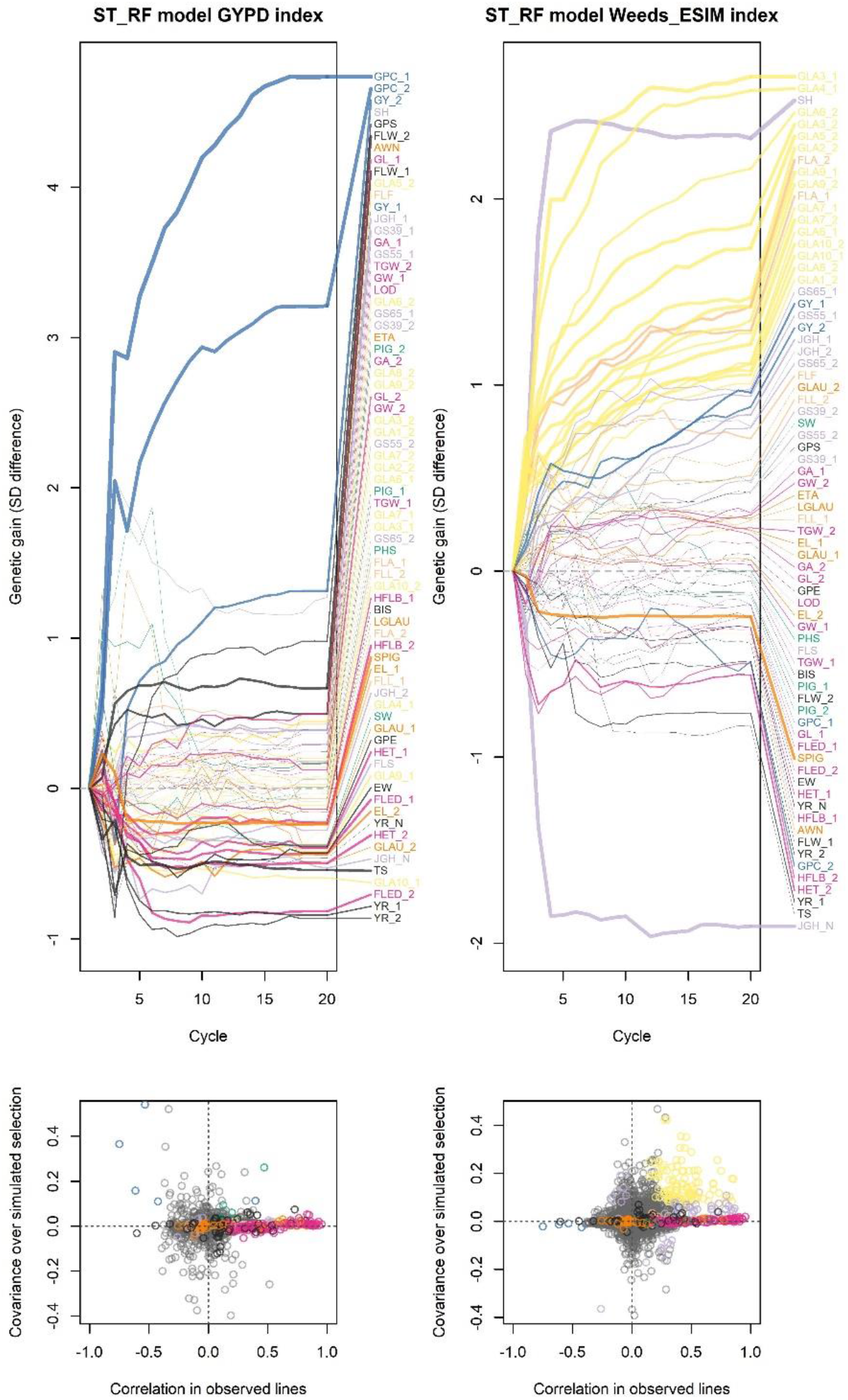
Simulated phenotypic response to selection on two selection indices. Upper plots indicate response of all traits under simulated recurrent selection for two multi-trait selection indices based on single-trait Random Forest (ST RF) genomic prediction models. GYPD = selection to increase both grain yield and protein (grain yield protein deviation); Weeds_ESIM = selection to increase yield as well as weed competitive traits whilst limiting change in plant height. Line colours relate to trait groups identified by hierarchical clustering and correlations shown in Figure 1. Line widths are relative to the t-test significance of each trait genetic gain from cycle generation 0 to 20 across all 20 simulation repeats. Lower plots compare the correlations between all pairs of traits in the original population and the covariance between trait pairs over time in the simulated population under selection. Points in upper right or lower left quadrants indicate both positive or negative correlation and covariance which demonstrates, suggesting complementary trait selection. Points in the lower right or upper left quadrants represent differing positive or negative correlation and covariance which demonstrates, suggesting antagonistic trait selection. Point colours indicate pairs of traits that are both in the same trait group following the colour scheme in Figure 1, while grey points indicate trait pairs from different groups.

Selection on multiple traits, which requires optimisation of multiple trait trade-offs, slowed the rate of genetic gain in grain yield for each of the selection indices: mean grain yield across both trial years increased when using both selection indices, but at a slower rate, particularly for grain yield in year 1 under GYPD selection, compared to when grain yield was selected for *per se* (**Figure 8**). However, in comparison to gain in grain yield in either one of the two years when selection was for grain yield in the other year, both GYPD and the yield + weed competition selection indices achieved generally comparable gains for yield whilst also increasing other favourable traits (**Figure 8**). These results show that antagonistic trait relationships are generally possible to optimise through appropriate selection. However, while this may slow genetic gain to some extent in the primary traits of interest, such as grain yield, this more realistically represents the balance of selection for multiple traits that occurs in breeding programmes.

**Figure 8.**
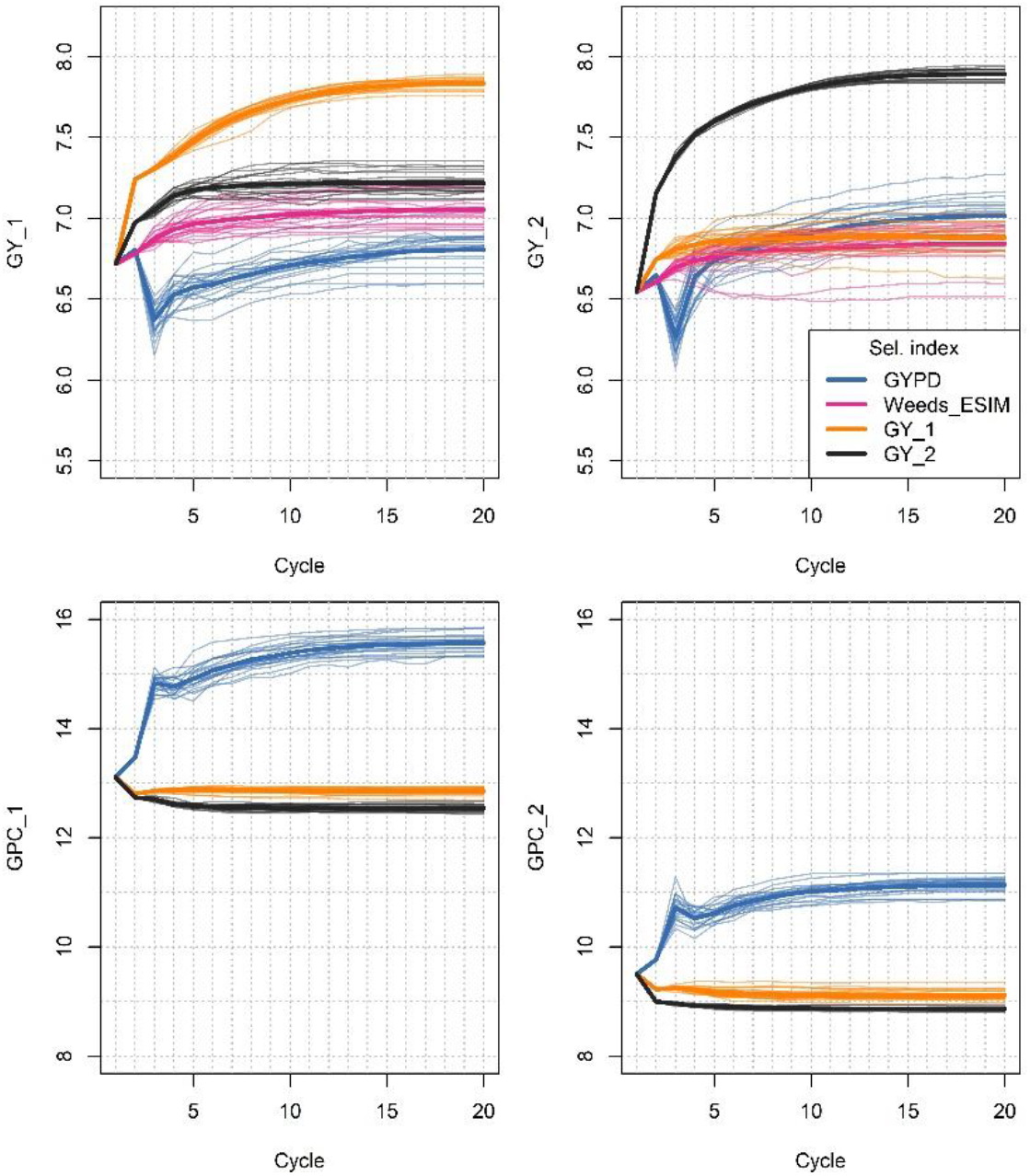
Simulated response to selection in grain yield (GY) measured in two trial years (_1 = year1, _2 = year2) under different selection indices based on single-trait Random Forest (ST RF) genomic prediction models. Narrow lines represent each of 20 simulation repeats while thicker lines represent the mean across all simulation repeats. GYPD = selection to increase both grain yield and protein (grain yield protein deviation); Weeds_ESIM = selection to increase yield as well as weed competitive traits whilst limiting change in plant height.

### 3.6 Different true genetic models affect long-term response to simulated selection

After comparing simulated response to different selection indices, we then tested how using different true genetic models that are based on the different genomic prediction models trained on the observed NDM population (LASSO versus RF; using either ST or MT approaches) affect phenotypic and genomic response to selection. Both RF and MT ensemble models were shown above to generally increase the prediction accuracy across traits (**Figure 3**), and here we show these predictions had a lower degree of shrinkage towards the mean of predicted trait values in comparison to the grain yield trait values of observed lines (**Figure 9a**). The MT RF models in fact had comparable variances in prediction values to the observed grain yield data, indicating their realistic prediction of phenotypic variation. While this would be expected from an overfitting model, cross validation with the observed data showed an increased accuracy of these models.

**Figure 9.**
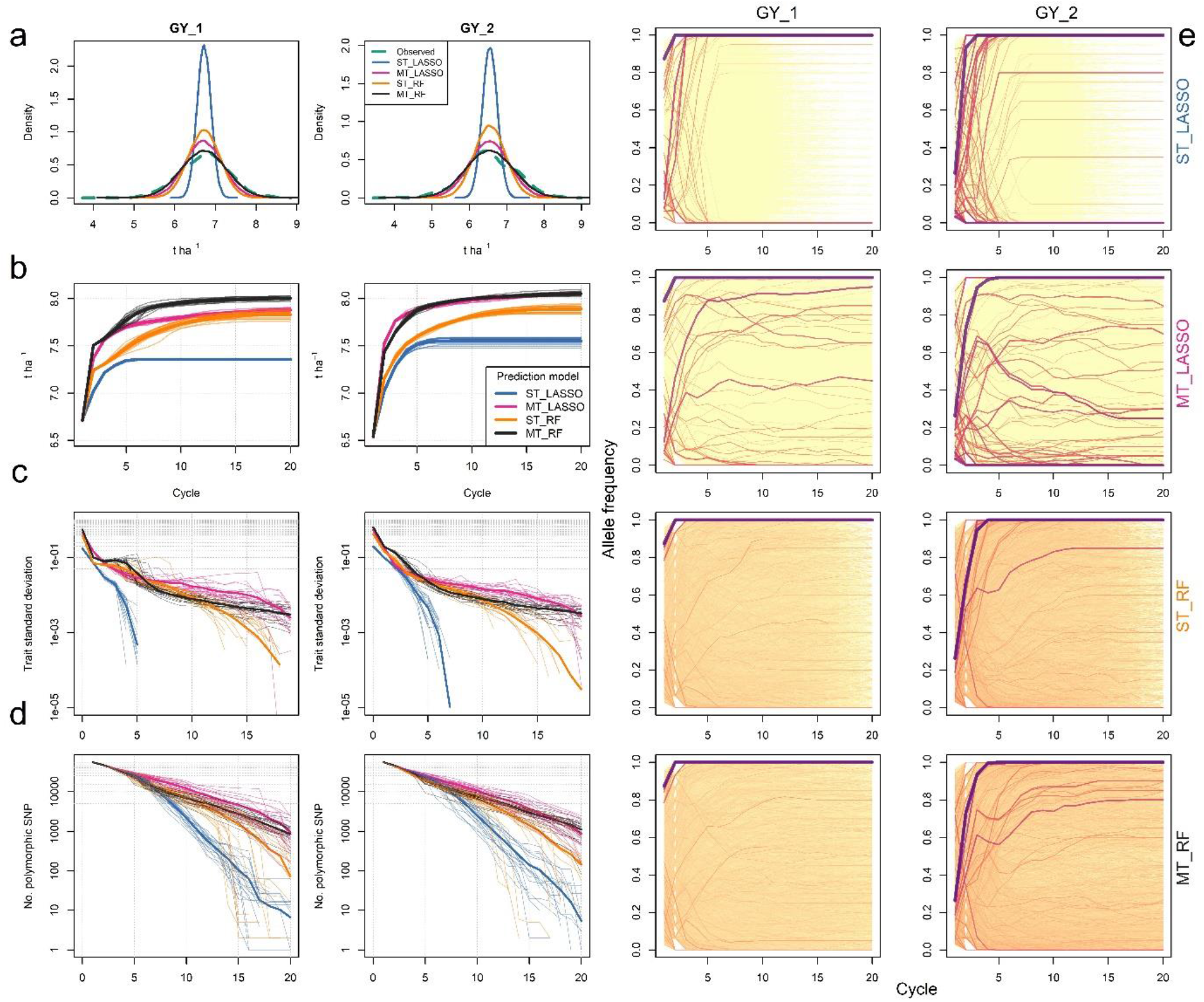
Simulated phenotypic and genetic response to selection with different genetic models. (**a**) Comparisons among distributions of observed and predicted trait values for two grain yield (GY) scenarios for different prediction models. LASSO and Random Forest (RF) genomic prediction models in conjunction with single-(ST) or multi-trait (MT) ensemble models are compared. (**b**) Rates of genetic gain in GY in each GY scenario when GY is directly selected under 20 cycles of simulated recurrent selection comparing different genomic prediction models. Comparisons in the rate of reductions in (**c**) phenotypic and (**d**) genetic variation in the NIAB Diverse MAGIC (NDM) population under recurrent selection comparing the same prediction models as above, and colour coded in the same way. Narrow lines represent each of 20 simulation repeats while thicker lines represent the mean across all simulation repeats. (**e**) Changes in mean allele frequency for all ~55,000 SNP markers across 20 simulation repeats for the NDM population under simulated selection for grain yield in two yield scenarios using the four models (ST LASSO, ST RF, MT LASSO, MT RF). Line widths and colour are proportional to SNP effect size in LASSO models and variable importance score for RF models.

Simulations with MT RF true genetic models tended to have the largest and longest increase of genetic gain over the course of simulated selection of grain yield (**Figure 9b**). Genetic gain in grain yield plateaued, at a relatively low level, after only around six cycles of selection using ST LASSO genetic model simulations but both cycle time to plateaux and plateaux height (maximum genetic gain) were both extended by either using a RF or MT model. This pattern of faster and higher genetic gain in RF or MT models was accompanied by the retention of higher phenotypic (**Figure 9c**) and genetic (**Figure 9d**) variance, particularly over long-term selection in the MT models. Almost all non-zero LASSO SNP effects were fixed after 8 cycles of selection in any simulation repeat for selection for grain yield in either year, limiting further genetic gain (**Figure 9**). Continued loss of genetic diversity once all genetic effects that affect phenotypic variance were fixed was down to genetic drift. Many of the SNP with highest variable importance in RF models were in common with the largest LASSO SNP effect coefficients, and the largest of these were fixed in the first few cycles of selection at a similar rate for both ST RF and ST LASSO (**Figure 9e**), where almost all of the ten SNPs with the largest LASSO effect or RF variable importance were fixed after five cycles of selection for both models. However, RF models included many more SNPs with non-zero importance (~20,000) than non-zero LASSO effects (61 and 87 for grain yield in years 1 and 2, respectively) and many more of these small or non-additive genetic effects in RF models remained polymorphic for longer (**Figure 9e**). For example, a significant proportion of these (14.3 and 13.8% for grain yield in years 1 and 2, respectively) remained polymorphic after 10 cycles of selection while genetic gain in yield still continued to increase (**Figure 9b**). This suggests that accumulation of the SNP effects, that were too small to be included in LASSO, or complex non-additive SNP by SNP epistatic genetic effects, made a large contribution to continued long-term genetic gain in RF models even after large effect QTL are fixed.

Furthermore, simply adding a MT second step to LASSO models to include indirect pleiotropic effects also increased and extended long-term genetic gain to a similar or greater extent to ST or MT RF models (**Figure 9b**). Using MT models, LASSO SNP effects were fixed at a much slower rate (**Figure 9e**) and phenotypic and genetic variance was maintained for much longer (**Figure 9c-d**), where on average 12% of the ten largest LASSO SNP effects for each single trait were polymorphic after 10 cycles of selection for across all simulation repeats with selection for grain yield in both years. This also suggests that the greater degree of pleiotropy present in MT models, which increased prediction accuracy for low accuracy LASSO models in particular (**Figure 3**), meant that the number of small effect loci involved in each trait was greatly increased. However, the number of indirect pleiotropic LASSO SNP effects across non-additive ensemble models could not be quantified. Selection could therefore act on more complex trait relationships driven by pleiotropy and/or linkage.

Linkage among antagonistic genetic effects could be shown to partly limit genetic gain. On average, only 0.8 and 5% of non-zero ST LASSO model SNP coefficients were negatively fixed resulting in an average of 0.28 and 2.15% loss of the maximum yield after 20 cycles of selection for yield in each year respectively. However, this was exacerbated in MT LASSO genetic models where 16.4 and 28.7% of ST LASSO SNP effects with negative effects on the trait under selection, were incorrectly fixed resulting in 14.1 and 23.5% loss of genetic gain. This further indicates insufficient recombination to completely decouple antagonistic linked QTL that were not directly involved in ST LASSO models for grain yield directly but pleiotropically linked through MT models.

## 4. Discussion

A complex structure of trait relationships that interact with environmental conditions were found to be involved in prediction of grain yield. Through simulation of recurrent selection within a genetically diverse highly recombined multi-founder wheat population, and based on observed genomic and phenotypic data, we tested several contrasting genetic models and quantitative genetic approaches to recurrent selection. We found that, in comparison to a simplifying LASSO genetic model where each trait was predicted directly from a minimal subset of markers with additive effects, prediction accuracies were increased both by using a multi-trait ensemble approach and Random Forest prediction models, which potentially incorporate pleiotropy and epistatic effects respectively. This was particularly so for complex traits with low prediction accuracy, and in simulations of recurrent selection these models also increased the rate and extent of long-term genetic gain, whilst maintaining phenotypic and genetic variance. Thus, genomic prediction models that include more complex genetic effects such as epistasis, and pleiotropy may better reflect how continued genetic gain is achieved through breeding.

### 4.1 The value of multi-trait models

We showed that modelling relationships among traits is valuable for increasing genomic prediction accuracy. Traditional multi-trait genomic prediction models consider the covariance structure of related traits across multiple environments and replicates and increase genomic prediction accuracy for cross-validation schemes when test fractions include partially phenotyped individuals in the test environment (Jia and Jannink, 2012). However, other studies often do not find an advantage to multi-trait models for untested genotypes in real datasets (Bhatta et al., 2020; Ward et al., 2019). We present results from multi-trait ensembles that integrate predictions of multiple traits into the same model (Van der Laan et al., 2007; He et al., 2016). These ensemble models consistently outperformed single trait models, while a contrasting approach using single vector decomposition of the multi-trait matrix performed poorly and more variably across traits. Although the increase in prediction accuracy was small for most traits, the advantage of multi-trait ensemble models was particularly great for traits that were poorly predicted by LASSO models, suggesting that ensemble methods efficiently incorporate additional information from large numbers of small pleiotropic genetic effects among related traits, which ST LASSO models would otherwise overlook when each trait is considered independently. Traits such as grain yield are polygenic and few genetic markers with large and consistent effects have been identified and applied in breeding (Bernardo, 2016). However, predictions of component traits of yield, many of which have simpler genetic architectures (Scott et al., 2021), can improve the ensemble prediction model for yield. We found that many highly correlated traits were used as covariates with high importance in multi-trait models. Furthermore, the covariate importance scores of traits in the ensemble models highlight physiological mechanisms for trait improvement and enable optimisation of antagonistic trait relationships (**Figure 4**). Where yield components correlate negatively with each other, the multi-trait ensemble model is able to optimise the interplay among these traits to increase the prediction accuracy of yield as the primary trait of interest. Similar to the approach taken by Powell et al. (2022) who modelled multiple systems biology development processes to bridge the gap between genotype to complex phenotype, we used multiple physiological traits in more agnostic models without defined crop growth parameters to aid prediction of the complex processes behind grain yield.

Inter-year environmental variation can modulate relationships between traits. We noted strongly contrasting weather conditions between the two trial years in which phenotypic data was collected (**Figure 2**). The covariate importances of traits for predicting yield changed with the year scenario being predicted, revealing some mechanisms controlling G×E for yield (**Figure 4**). For example, growth stage phenotypes were more important covariates in year 1. Similarly, trade-offs in plant size and earliness likely maintain polygenic trait variation due to varying environmental pressures in the wild plant *Mimulus guttatus* (Troth et al., 2018). In breeding, any single strategy to achieve high yield may be hampered by unpredictable year-to-year environmental variation, and thus limit response to selection and reduction in genetic variance. While our simulations of future genetic gain cannot account for unmeasured environments in the future, commercial wheat breeders often take this into account and make selections of promising lines with a diversity of phenological or plant height traits to ensure adaptive potential.

### 4.2 The potential to optimise trait trade-offs that conventional breeding has neglected

Using multi-trait data from a MAGIC population that controls for confounding effects of population structure (Scott et al., 2020), we found that pleiotropy and/or tight genetic linkage are significant causes of correlated trait responses to selection. These data also shed light on the combination of traits that would be required to be co-selected or optimised to achieve continuous gains in grain yield as a primary trait under selection. Furthermore, we find antagonistic trade-offs among traits that have been problematic for wheat crop improvement. We suggest that historic enhancement of grain yield by breeders at the cost of key traits such as weed competitive ability, or grain protein content, has been due to the over-riding value placed on grain yield as a primary selection criterion during variety testing, as well as market pressures, leaving little scope for compromise with other traits. Integration of novel trait variation to optimise these trade-offs within an elite wheat genepool, which has been under such strong directional selection, would therefore be difficult. However, simulations presented here show that with appropriate selection indices, genetic gain in both yield and other valuable but negatively correlated traits was possible to some extent.

### 4.3 The value of complex genomic prediction models for continued genetic gain

Simulating selection using prediction models with contrasting genetic architectures (i.e., in terms of additive or epistatic and direct or pleiotropic genetic effects) had major impacts on the outcomes of recurrent selection. The greater complexity of these models both increased cross-validated prediction accuracy of complex traits in the observed population and extended the accuracy of genomic predictions in simulations of recurrent genomic selection. Furthermore, simulations of phenotypic selection assuming a complex genetic model demonstrated accelerated and extended potential for genetic gain while maintaining genetic and phenotypic variance. The role of non-additive genetic effects has been demonstrated elsewhere to preserve genetic variance over long-term selection in simulated populations (Wientjes et al., 2021). Wang et al. (2004) used simulations of selection within the CIMMYT wheat breeding programme to compare genetic models, finding that inclusion of epistasis in genetic models greatly reduced the rate that additive genetic variance is lost due to selection. Although the role of epistasis is thought to contribute little to overall genetic variance, at least in outbred populations (Hill and Mäki-Tanila, 2015), evolutionary theory supports these observations in crop breeding; selection can enable conversion of epistatic to additive genetic effects, allowing hidden or cryptic genetic variation to then be unlocked (Carlborg et al., 2006; Hill, 2017). The limits to trait variation in our study are likely underestimates because as allele frequencies shift and trait genetic architectures evolve under selection new additive genetic effects would be unlocked for selection which cannot be modelled or predicted in the observed population. Supporting this, we found that prediction models soon become out of date and suffer loss of prediction accuracy, particularly for simple additive prediction models (LASSO), when target genotypes become more distantly related to the training set (Edwards et al., 2019). However, in realistic scenarios of a wheat breeding programme practicing genomic selection, the training model is continually updated with data from advanced breeding line testing which would enable more linear continued genetic gain. Continuous novel mutations may also play an important role in regenerating genetic variation and extending limits to long-term selection in large populations (Hill, 1982) but were not considered in simulations reported here. Pre-breeding programmes can also introduce novel genetic diversity from the primary, secondary and tertiary wheat genepool (Balfourier et al., 2019). Nevertheless, the population we study is representative of diverse north-west European wheats across 70 years. We found that additive variation included in minimal LASSO prediction models was quickly depleted during simulated selection. We propose that pleiotropic and epistatic genetic effects and G×E interactions have played a major role in maintaining wheat genetic diversity despite strong selection and will be particularly important for applied genomic selection of elite varieties in already highly selected breeding populations.

### 4.4 Potential for applied crop breeding

MAGIC populations have proven valuable resources for direct generation of commercial varieties of some less intensively bred crops than wheat (Scott et al., 2020). In simulated breeding programmes, Bernardo (2021) suggested that multi-parent crossing schemes may be valuable for maintaining genetic diversity. However, the diverse MAGIC wheat population described here is unlikely to generate commercially competitive varieties due to the broad genetic basis and historic founders. Instead, this MAGIC population samples and recombines genetic diversity across 70 years and can therefore be considered a microcosm of past and future selective breeding. In this context, our simulations rerun alternate histories to test different selection models and approaches and reveal physiological and genetic mechanisms for future breeding. We suggest that this approach, including multi-trait ensembles, could be further integrated with environmental information to inform crop models (Cooper et al., 2021; Stöckle and Kemanian, 2020). Considering that traditional wheat breeding programme cycles generally extend over at least five years, our simulations of twenty cycles of recurrent selection represent an equivalent of over one hundred years of traditional wheat breeding (albeit it with no further input from genotypes outside of the 16 founders from which the MAGIC population was constructed). Current wheat breeding programmes are also likely to be at a point towards the later stages of selection simulations presented here where the majority of large effect QTL are either fixed or well accounted for. Further genetic gain in current breeding programmes will therefore likely be achieved through optimisation of small and complex genetic effects (Gorjanc et al., 2018).

Through selection, breeders appear to often maintain linkage disequilibrium across unexpectedly large genomic regions because they contain several beneficial alleles (Brinton et al., 2020; Fradgley et al., 2019), which could also interact epistatically. Therefore, prediction models that capture relevant haplotype blocks and the most recently unlocked epistatic effects will be increasingly important for forward prediction of high performing breeding lines. Comparisons between patterns of linkage disequilibrium in commercial selected varieties and in simulations presented here would validate this process and uncover valuable sites for further marker assisted selection. Our simulations suggest that beneficial variation is often lost during breeding. This diverse MAGIC population is a reservoir for such this genetic diversity and the phenotypic and genomic data allow beneficial alleles to be identified. Through targeted rapid recurrent selection and breeding technologies that reduce generation time (Watson et al., 2018; Cha et al., 2021), this population could therefore be used to provide useful pre-breeding material for commercial breeding programmes to deliver accelerated and continued genetic gain. Optimum contribution selection can also be applied to maximise long-term genetic gain by maintaining genetic variance for selection at later generations.

### 4.5 Conclusions

In summary, we demonstrated the value of multi-trait ensemble models for genomic prediction of complex traits and simulated recurrent selection using these genetic models based empirically on an extensively genotyped and phenotyped NDM population. We consider this a microcosm of wider wheat breeding programmes working with the wider pool of wheat germplasm so that our results provide insights into the trends and mechanisms by which the considerable progress and genetic gain has been made in modern wheat breeding without apparent genetic diversity loss. These findings highlight the importance of models and approaches that take into account these mechanisms to maximize further genetic gain in the future.

## 5. Acknowledgements

This work was funded by Biotechnology and Biological Sciences Research Council (BBSRC) grants BB/M011666/1 awarded to JC, BB/M011585/1 to RM and BB/M011194/1 as part of the PhD research of NF. We also thank Dr Stephanie Swarbreck for useful comments on the manuscript.

## 6. Author contributions

NF conceived of and undertook the analyses. JC and RM were awarded project funding. NF wrote the first draft of the manuscript and all authors reviewed, edited, and approved the final manuscript.

## 7. Data availability

Phenotypic and genotypic data used in this study were as presented by Scott et al. (2021) and can be freely accessed at http://mtweb.cs.ucl.ac.uk/mus/www/MAGICdiverse/.

## Notes

http://mtweb.cs.ucl.ac.uk/mus/www/MAGICdiverse/

### Competing Interest Statement

The authors have declared no competing interest.

