## Supplementary figures and images for "Multi-trait ensemble genomic prediction and simulations of recurrent selection highlight importance of complex trait genetic architecture in long-term genetic gains in wheat"

### Supplementary Figure S1

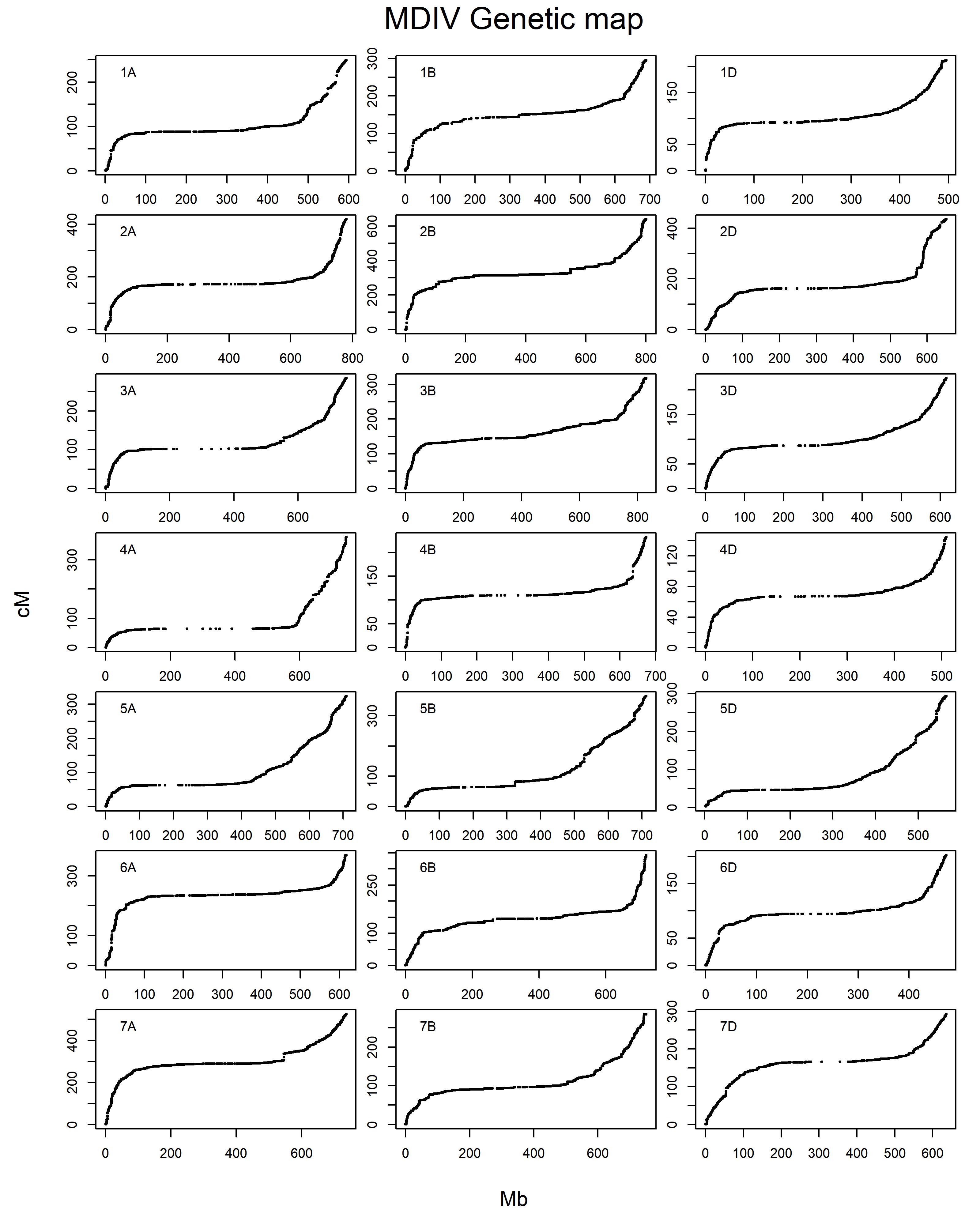
